# Experimental evidence for the impact of soil viruses on carbon cycling during surface plant litter decomposition

**DOI:** 10.1101/2021.09.29.462240

**Authors:** Michaeline B.N. Albright, La Verne Gallegos-Graves, Kelli L. Feeser, Kyana Montoya, Joanne B. Emerson, Migun Shakya, John Dunbar

## Abstract

To date, the potential impact of viral communities on biogeochemical cycles in soil has largely been inferred from indirect evidence, such as virus-driven changes in microbial abundances, viral auxiliary metabolic genes, and correlations with soil physiochemical properties. To more directly test the impact of soil viruses on carbon cycling during plant litter decomposition, we added concentrated viral community suspensions to complex litter decomposer communities in 40-day microcosm experiments. Microbial communities from two New Mexico alpine soils, Pajarito (PJ) and Santa Fe (SF), were inoculated onto grass litter on sand, and three treatments were applied in triplicate to each set of microcosms: addition of buffer (no added virus), addition of live virus (+virus), or killed virus (+killed-virus) fractions extracted from the same soil. Significant differences in respiration were observed between the +virus and +killed-virus treatments in the PJ, but not the SF microcosms. Bacterial and fungal community composition differed significantly by treatment in both PJ and SF microcosms. Combining data across both soils, viral addition altered links between bacterial and fungal diversity, dissolved organic carbon and total nitrogen. Overall, we demonstrate that increasing viral pressure in complex microbial communities can impact terrestrial biogeochemical cycling but is context-dependent.

## Introduction

Viruses that infect microbial hosts have a major impact on their immediate host, but can also influence larger scale environmental processes. Viruses account for an estimated 20-40% of bacterial mortality in the oceans and are believed to be a major driver in marine carbon (C) cycling (1, 2). Viral infection likely impacts C and nutrient cycling in aquatic systems (3), and several pathways related to broad scale impacts on the marine carbon cycle have been proposed (4). The ‘viral shunt’ emphasizes the recycling on dissolved organic matter (DOM), ultimately fueling CO2 release into the atmosphere by heterotrophic microbes (5), while the ‘viral shuttle’ emphasizes viral driven organic particle aggregation that favors carbon export to the deep ocean (6).

As in marine systems, recent research suggests that viral mediated processes also impact terrestrial C and nutrient cycling (7–10). However, the magnitude and nature of these impacts is unknown. Differences between marine and terrestrial ecosystems that may influence the impact of viral mediated C cycling include differences in microbe and virus turnover times and the environmental spatial structure and heterogeneity (10, 11). Efforts to characterize viral diversity in soils and their impacts on biogeochemical cycling are rapidly increasing and have focused on exploiting previously sequenced metagenomic (12) and metatranscriptomic (8) datasets as well as creating novel viromics datasets (13). To date, the impact of the soil virome on biogeochemical cycling has been inferred from viral driven changes in microbial populations (e.g. predator-prey cycles) (e.g.(14, 15)) and presence of auxiliary metabolic genes which may be impacting biogeochemical cycling (7). Direct evidence of viral community-mediated alterations in C cycling is lacking, but there is recent experimental evidence for soil viruses affecting nitrogen cycling (16). Given that terrestrial systems contribute approximately 50% of C efflux to the atmosphere, understanding the impact of soil viruses may be critical to enable better modeling of greenhouse gas emissions under climate change scenarios (17).

Changes in the composition of microbial communities are increasingly recognized as a factor that can drive substantial variation in soil carbon cycling (18, 19). Viral predation in litter decomposer communities is a mechanism that can not only alter community composition but also directly affect carbon flow. To assess the relevance of viral predation, we manipulated virus abundance in microcosm experiments. We inoculated two distinct soil microbial communities spiked with a live or killed viral community concentrate extracted from the same source soil into microcosms containing plant litter and sand substrate; a no virus treatment where buffer was added was used as an additional control (Figure 1, Supplemental Methods). We measured respiration in the microcosms over 40 days and measured dissolved organic carbon (DOC), total nitrogen (TN) abundance, and bacterial and fungal composition at the 40-day endpoint. We hypothesized that increasing the quantity of virus would alter microbially driven C cycling during plant litter decomposition by changing microbial community dynamics.

**Figure 1.**
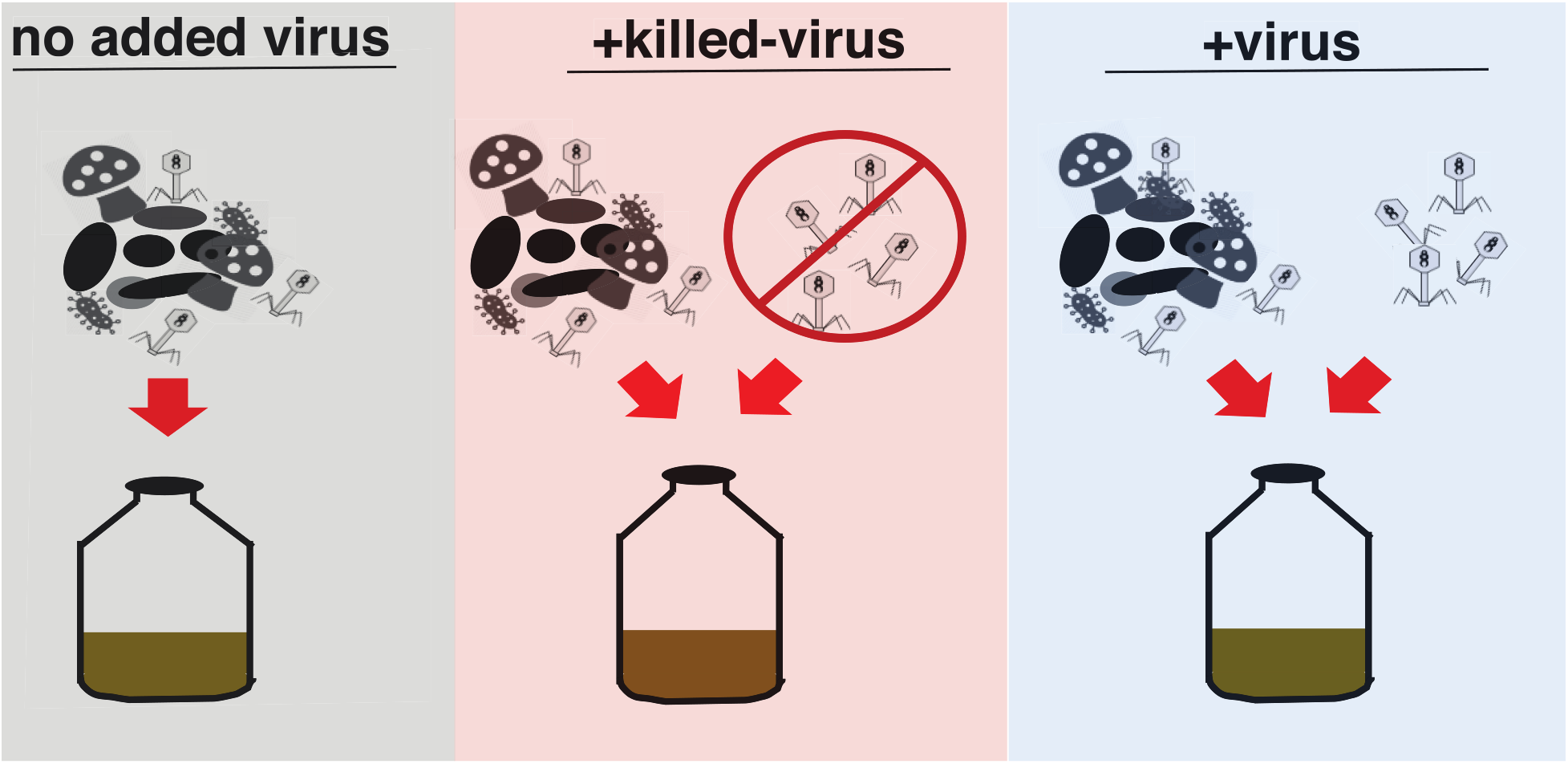
Experimental design. Treatments for each soil type included 1) whole microbial community (no added virus), 2) whole microbial community and killed virus concentrate (+killed-virus), and 3) whole microbial community and virus concentrate (+virus). Whole communities and viral concentrates were mixed together and added to microcosms containing sterilized plant litter on sand.

## Results and Discussion

### Virus extraction efficiency and magnitude of virus additions

We used lambda phage additions to soils to test the efficiency of our virus extraction protocol. Based on plaque assays, our soil virus extraction procedure recovered the lambda phage from soil with at least 50% efficiency (Figure S1). Given this, our virus addition treatment in microcosms was estimated to increase virus abundance over 15-fold. Estimates of the average viral abundances in soil are 1.5 × 10^7^ viruses per gram of soil (20), so this addition would amount to at least 2.8 × 10^8^ extra viruses. Of course, recovery of soil viruses may be slightly lower since extraction efficiency can vary among viruses owing to differential binding to filters during the extraction protocol (21). While our virus addition is artificial, large increases in virus abundance in ecosystems are expected when viral lytic cycles are triggered (22).

### Viral additions impact ecosystem processes

The impact of virus addition was assessed by comparing the +virus treatment to two controls: no added virus and +killed-virus (Figure 1). Based on visual inspection, our soil virus extracts contained additional organic matter (OM), so the +virus treatment inevitably boosted nutrient abundance in addition to viral abundance in the microcosms. The + killed-virus treatment was used to assess the impact of the nutrient addition. Initially, over the first eight days post inoculation, respiration increased in both the +virus and +killed-virus treatment compared to the control treatment in both the PJ and SF soil (Figure 2A), likely driven by the OM addition. However, in the PJ microcosms by day 14, respiration was significantly higher in the +virus treatment than the +killed-virus treatment and differences between these treatments increased to 30% by day 41 (Figure 2A). By contrast, in the SF microcosms after the initial surge in respiration in the +virus and +killed-virus compared to the no added virus treatment, by day 14 respiration in all samples was similar and this persisted (Figure 2A). Our hypothesis was partially supported; with one microbial community increasing virus quantity altered respiration, while with the other microbial community it did not. We posit that differences in the abundance and or diversity of viruses in the original soils and thus viral concentrates may have led to these microbial community specific responses. Given the high spatial variation observed in viral communities in soils in a previous study (13), differences in the spatial patterns of virus may impact soil C cycling.

**Figure 2.**
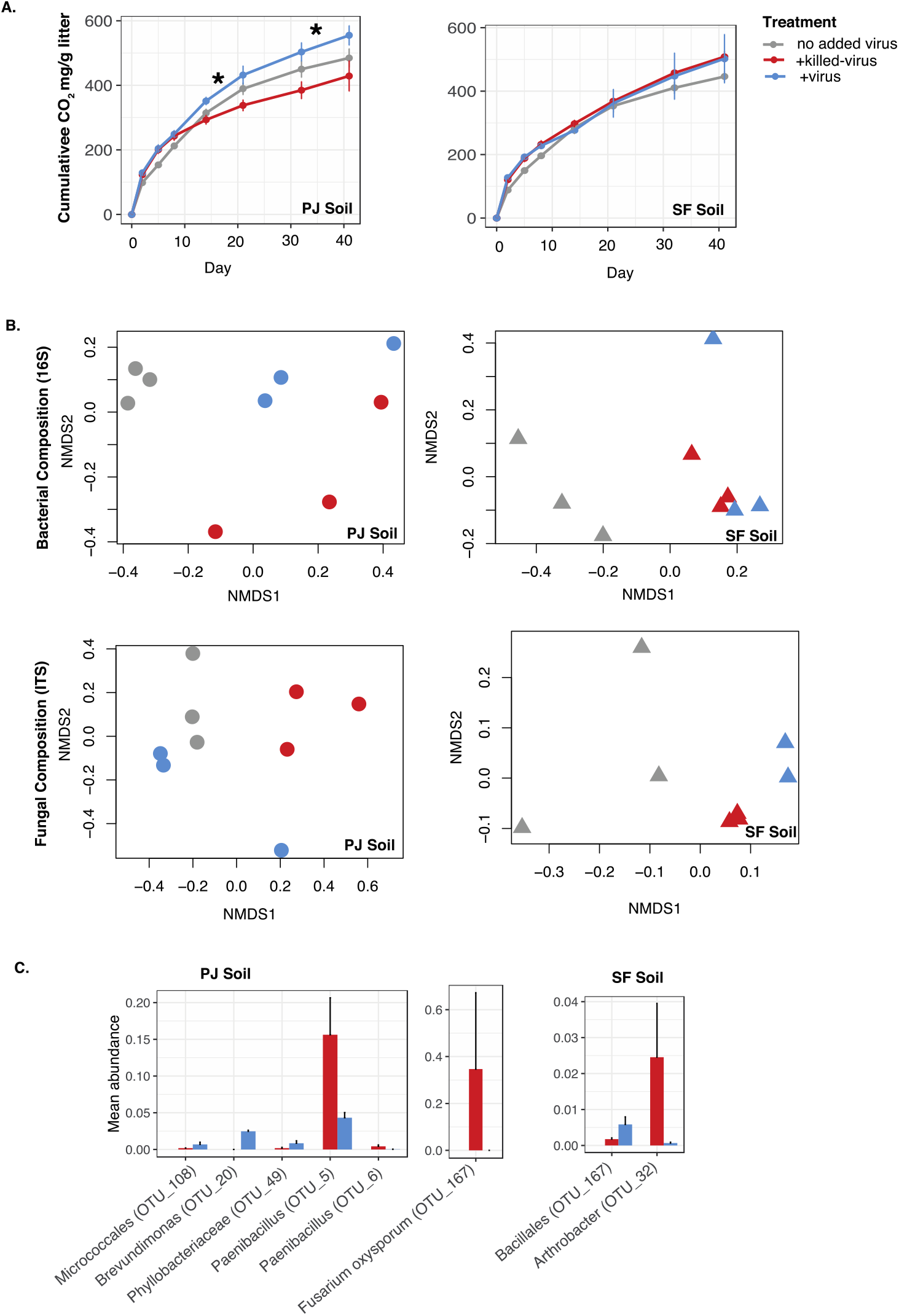
Viral addition impacts ecosystem functioning and microbial composition. **A)** Cumulative respiration across the treatments over 41 days for PJ (left) and SF (right) inoculated microcosms. Significant differences between +virus and +virus-killed treatments are shown (*) **B)** Bacterial (top) and fungal (bottom) composition in PJ (left) and SF (right) inoculated microcosms **C)** Mean relative abundance of taxa showing significant differences across the +killed-virus and +virus treatments identified through indicator species analysis (27). (See Figure S6 for an alternative analysis where differentially abundant taxa were identified between +killed-virus and +virus treatment using a DESeq2 analysis (28))

DOC abundance was lower in the PJ compared to the SF microcosms (Figure S2). While this trend was not statistically significant, the PJ +virus treatment generally had lower DOC abundance compared to the +killed-virus treatment, while for the SF microcosms we saw the opposite trend (Figure S2). We used a DOC mineral binding assay to assess changes in the DOC quality between the soil types in +virus versus +killed-virus treatments. Our results suggest that while autoclaving the inoculum samples (+killed-virus) changed the initial DOC quality compared to the non-autoclaved inoculum samples (+virus), these effects did not persist as there was no significant difference in DOC quality between treatments at 41 days (Figure S3).

### Viral addition impacts bacterial and fungal composition

As we hypothesized, virus additions significantly altered microbial community composition. We observed significant differences across treatment groups (no added virus, +virus, +killed-virus) for bacterial (F_2,8_ = 4.44, p=0.02, r=0.58; F_2,8_ = 3.11, p=0.02, r=0.51) and fungal composition (F_2,8_ = 2.06, p=0.02 r=0.40; F_2,7_ = 3.57, p=0.03, r=0.59) in the PJ and SF microcosms, respectively (Figure 2B, Figure S5). In particular, two OTUs in the bacterial genus *Paenibacillus* and one OTU belonging to the fungal species *Fusarium oxysporum* increased in relative abundance in the PJ microcosms, and one *Arthrobacter* OTU increased in relative abundance in the SF microcosms in the +killed-virus compared to the +virus treatment (Figure 2C). We posit that these taxa were impacted by predation in the +virus treatment, reducing their relative abundance. Increased viral predation on a few key taxa may have cascading impacts on overall microbial community assembly and successional processes during plant litter decomposition. Overall, the richness and Shannon diversity of bacteria and fungi in the +virus and +killed-virus treatments were reduced compared to the control treatment in both PJ and SF microcosms (Figure S4, Figure S6), likely due to nutrient addition, primarily OM residuals from the viral concentration and extraction from soil. Nutrient addition often reduces microbial richness (23). In the PJ microcosms we saw a trend of increased bacterial and decreased fungal richness and Shannon diversity in the +virus compared to the +killed-virus and the opposite trend in the SF microcosms, but it was not significant in either case (Figure S4).

### Viral addition alter links between microbial community traits and ecosystem processes

To test the impact of viral addition on links between the microbial community and ecosystem processes, we aggregated carbon, nutrient, and microbial community trait (bacterial and fungal diversity) metrics across both soil types for each of the three treatments. This analysis provided additional support of our hypothesis, where viral addition altered the relationship between carbon and nutrients and microbial community traits (Figure 3, Figure S7). In our two control treatments we saw positive correlations between bacterial and fungal diversity, which were lost in our +virus treatment. While, in the +virus treatment we saw a negative correlation between DOC and bacterial diversity, which was not observed in the two control treatments. Additionally, in the +virus treatment we observed multiple strong correlations between bulk and bound DOC and TN, which may point to viral driven signature in the dissolved OM(24). Finally, it is interesting to note that the +virus outcomes are not simply an additive result of nutrient addition and predation, another illustration of context dependence.

**Figure 3.**
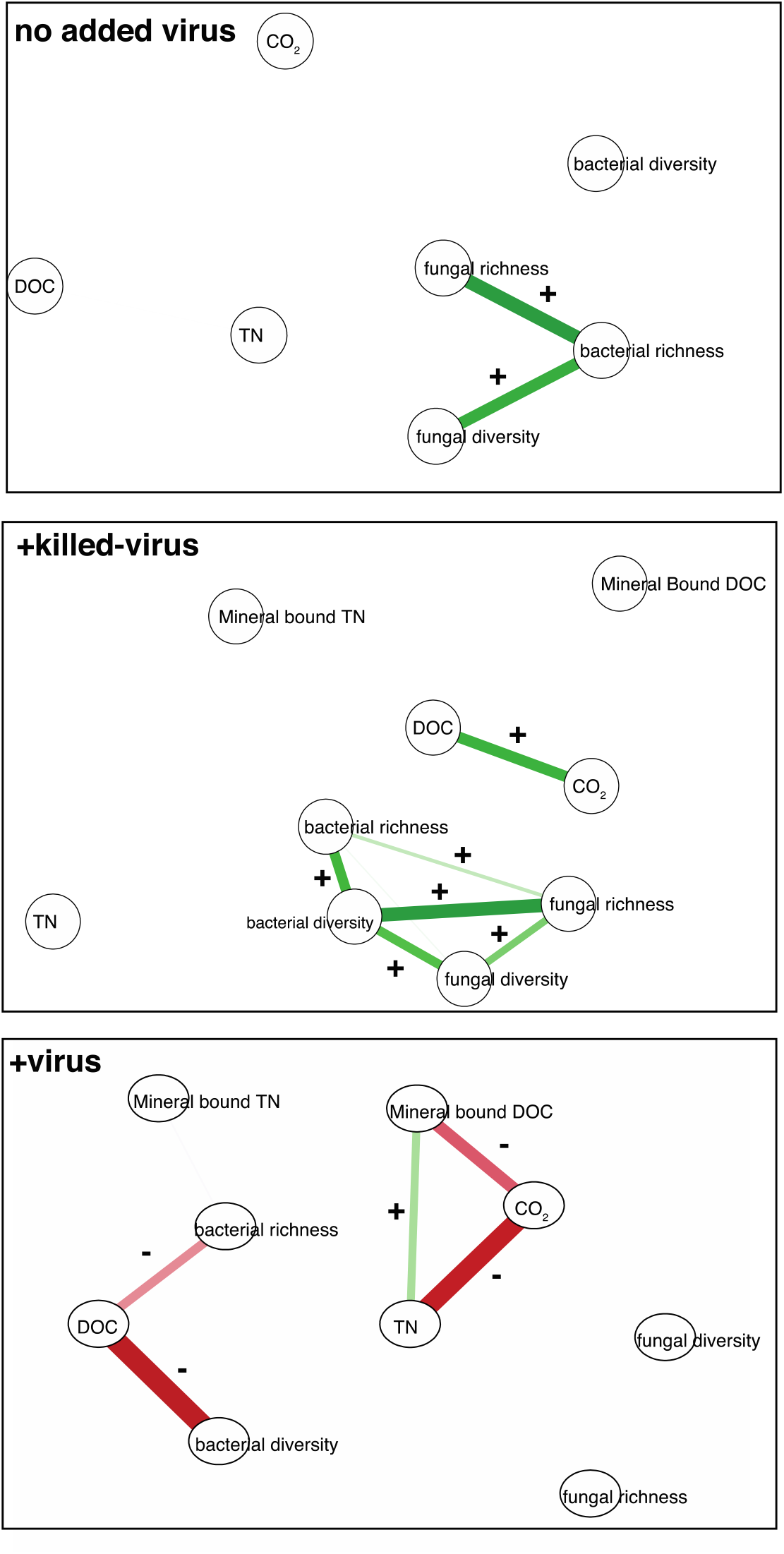
Viral addition alters correlations between microbial community traits and ecosystem processes. Spearman correlations (p>0.05) between bacterial and fungal richness and diversity and carbon and nutrient measurements are shown. (See Figure S7 for a correlation heatmap with additional information about correlations across all variables).

## Conclusions

While previous observational studies have shown correlations between viral abundance and carbon compounds (2) and used meta-omics to infer functional impacts of viruses on C cycling (3, 7, 9, 25, 26), here we experimentally demonstrate the impacts of soil viruses on carbon cycling by manipulating virus abundance in complex microbial communities. Our results show that increases in soil virus abundance can impact carbon and nutrient cycling in terrestrial systems likely by altering microbial community dynamics. However, the magnitude of these effects depends on factors such as community composition and nutrient availability. Future research is needed to delve into virus-host spatial and temporal dynamics in soils, where the physical structure may change dynamics compared to a more well mixed marine ecosystem.

## Acknowledgements

This work was supported by a Los Alamos National Laboratory, Laboratory Directed Research and Development grant 20200252ER to M.B.N.A., M.S. and J.D. and by SFA grant 2018LANLF255 from the U.S. Department of Energy Office of Biological and Environmental Research to J.D. We thank Katrine L. Whiteson for helpful conversations regarding the data.

## Author contributions

MBNA and JD designed the study with input from MS and JBE. MBNA, LVGG, and KM performed experiments. MBNA performed data analysis and wrote the first draft of the manuscript. All authors contributed to manuscript editing.

