## Supplementary Figures for "Experimental evidence for the impact of soil viruses on carbon cycling during surface plant litter decomposition"

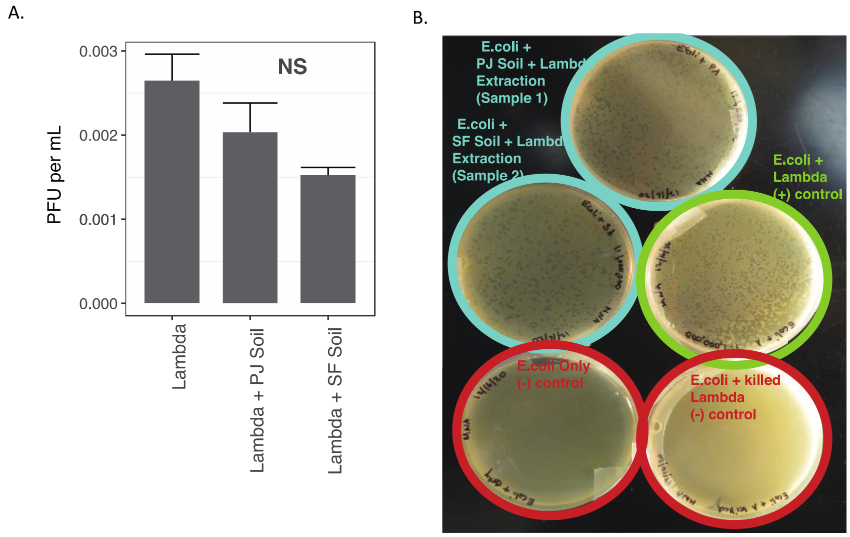


**Figure S1.** Phage extraction efficiency. A) Amount of lamda phage added to soils (left bar), and the amount of lambda phage that was extracted from the PJ and SF soils. B) Photo of plaque assays showing the *E.coli* + lambda positive control (green), the *E.coli* + lambda extractions from soils (light blue) and the *E.coli* only and *E.coli* + killed lambda negative controls (red).

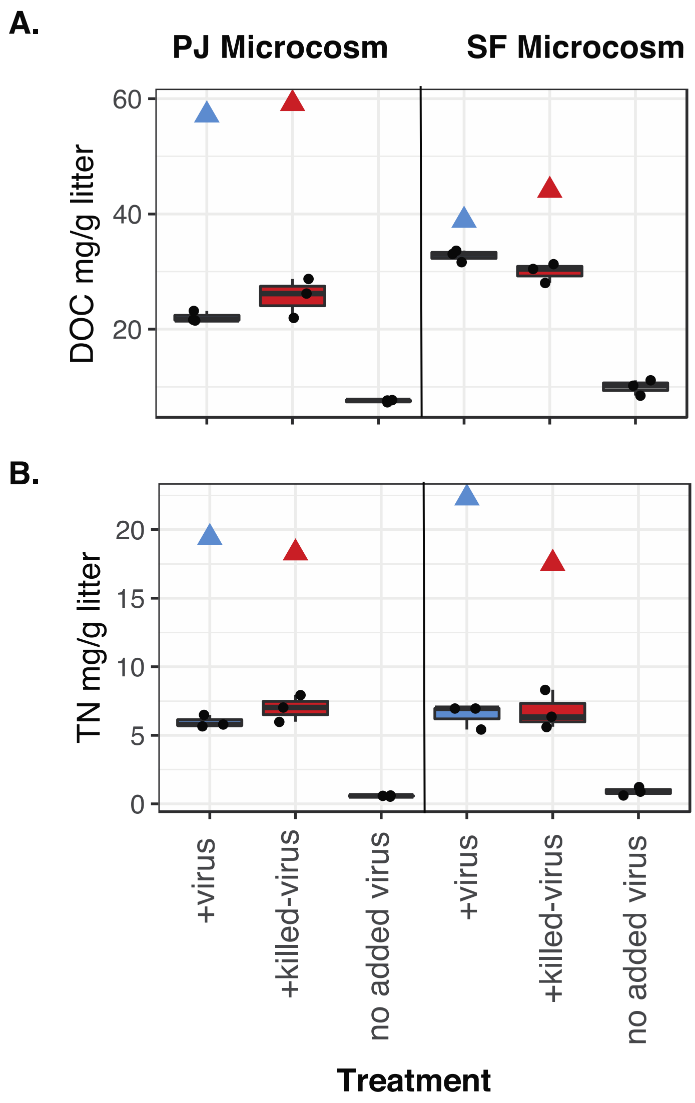


**Figure S2.** Dissolved organic carbon (DOC) and total nitrogen (TN) in microcosms inoculated with the PJ soil (left) and SF soil (right) after 40 days grouped by treatment, +virus (blue), +killed-virus (red), control (gray). Triangles show the concentration of DOC and TN added in the inoculum for the +virus and +virus-killed treatments.

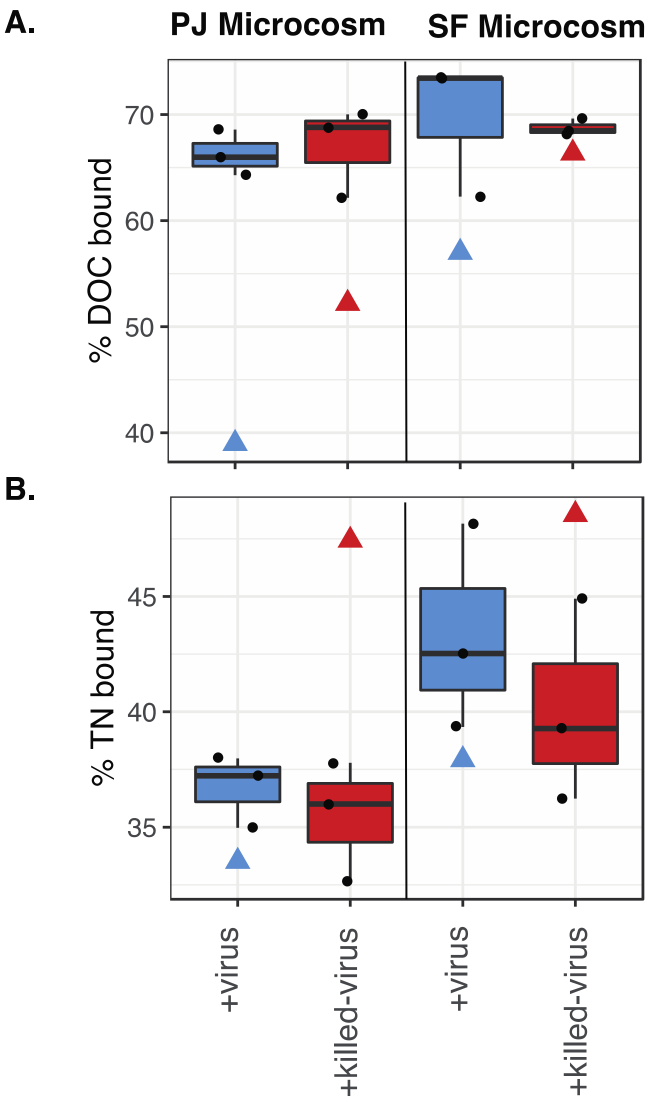


**Figure S3.** Percent of dissolved organic carbon (DOC) and total nitrogen (TN) that binds to aluminum oxide from the microcosms inoculated with the PJ soil (left) and SF soil (right). DOM was obtained from microcosms after 40 days of *blue grama* litter decomposition. Triangles show the DOM binding potential of the inoculum for the +virus and +virus-killed treatments.

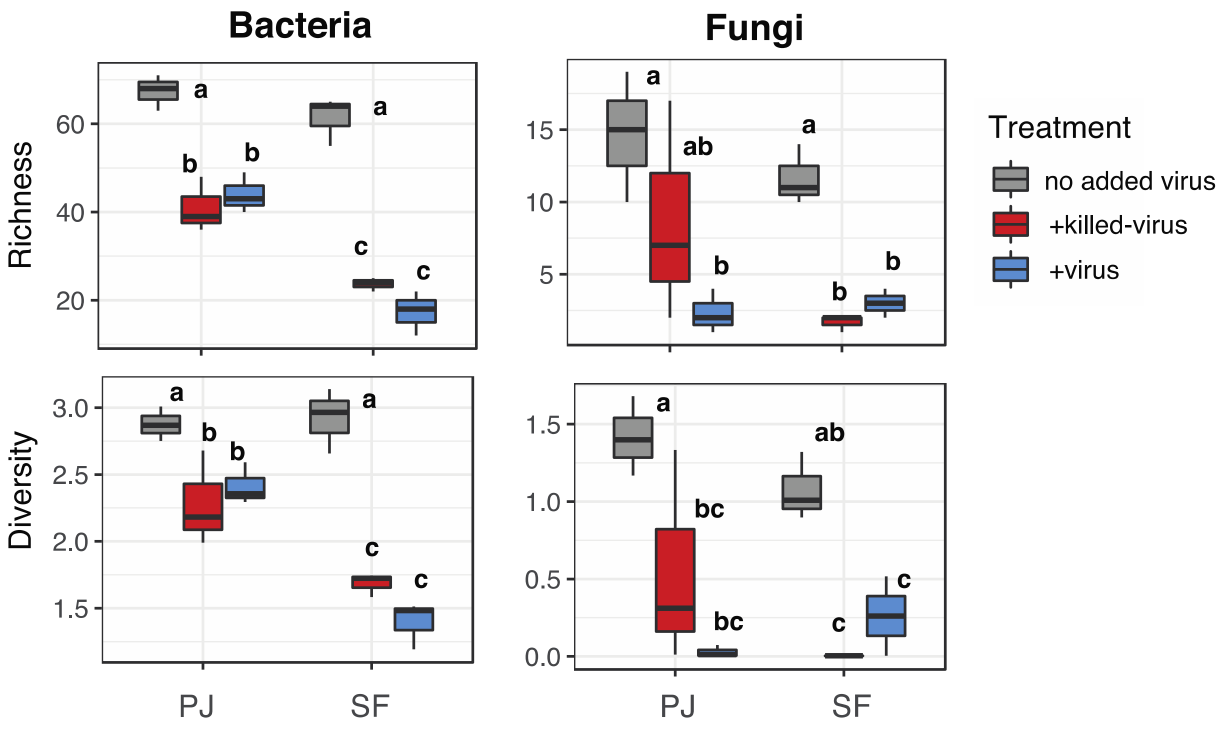


**Figure S4.** Bacterial and fungal richness and Shannon diversity across treatments for the PJ and SF microcosms. Letters indicate significant differences across treatments and soil inoculum (Tukey hsd posthoc test).

A.

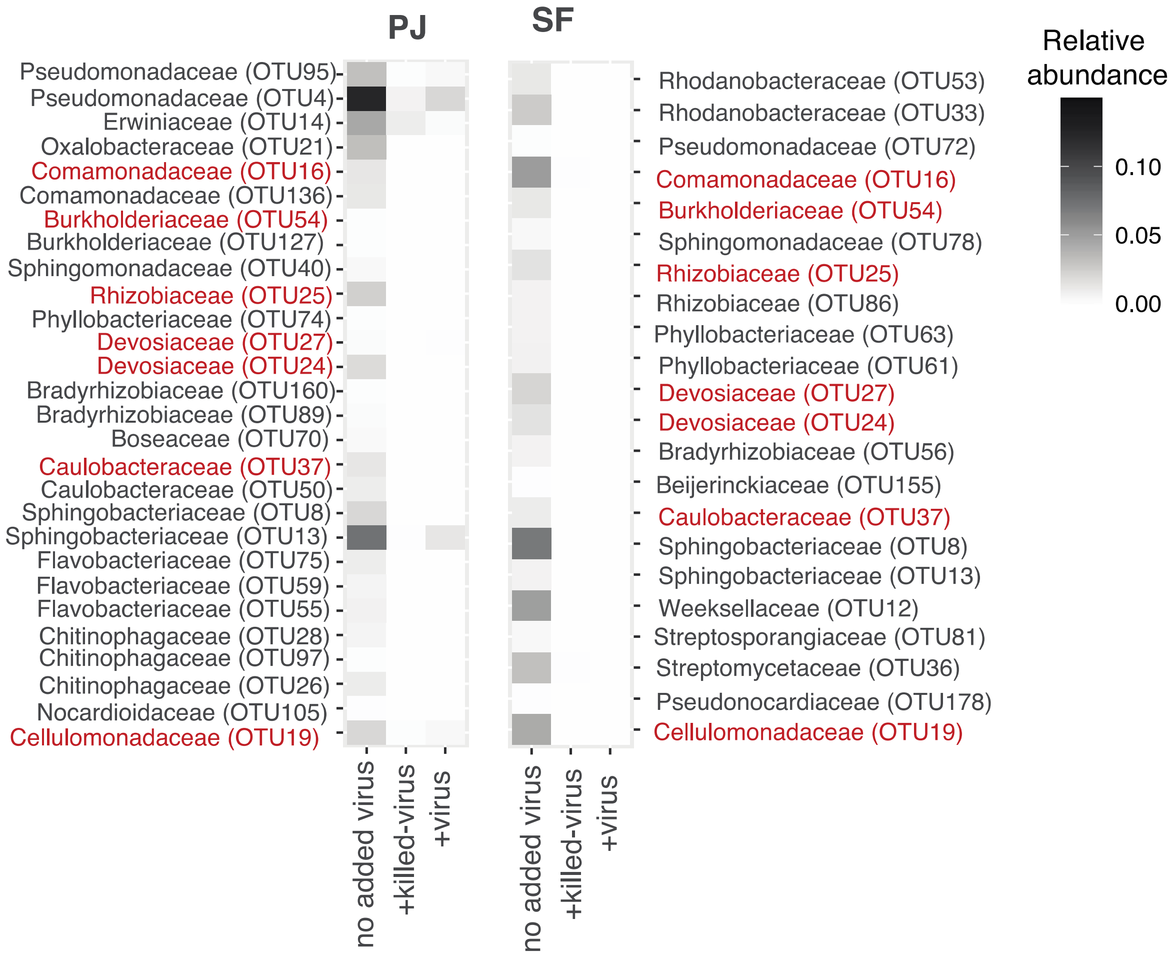


B.

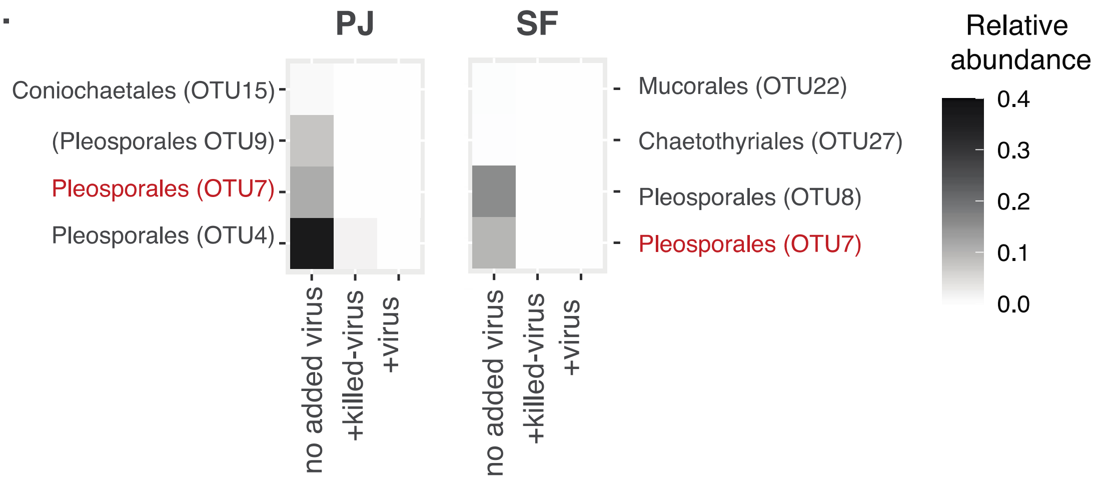


**Figure S5.** Mean relative abundance of A) bacterial and B) fungal taxa showing significant differences between the no added virus compared to the +killed-virus and +virus treatments identified through indicator species analysis. Taxa that were more abundant in the control treatment in both PJ and SF microcosms are highlighted in red.

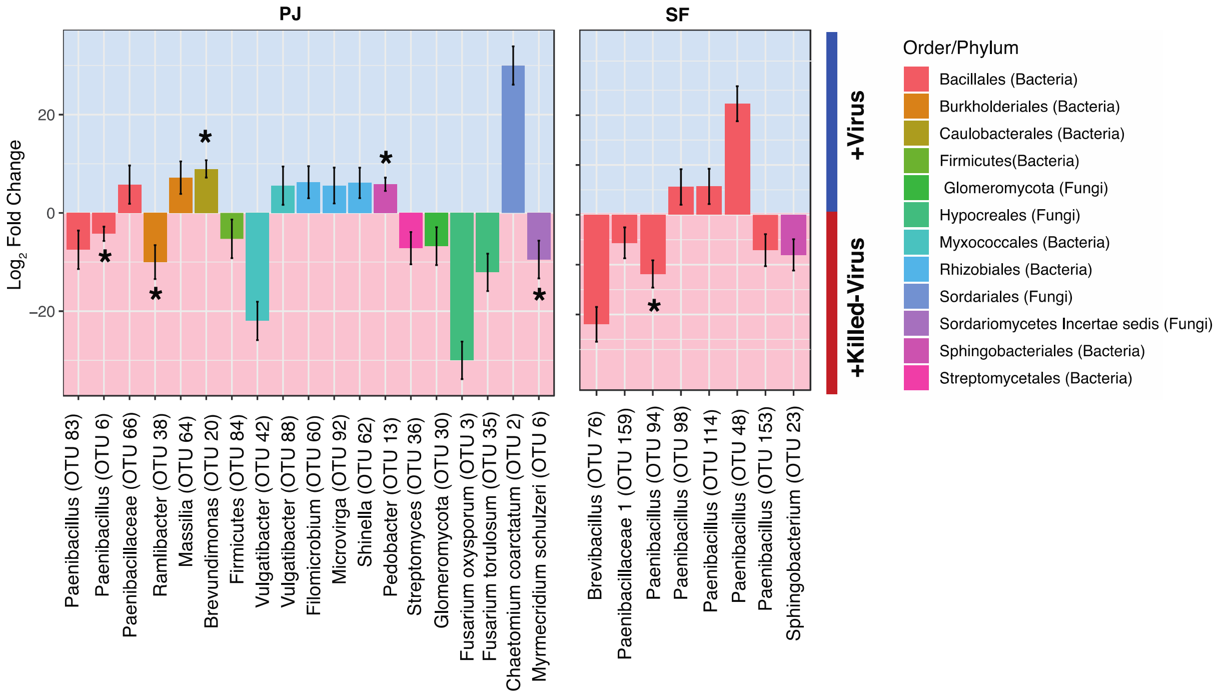


**Figure S6.** Differences in bacterial and fungal composition of +virus versus +killed-virus versus samples in PJ and SF microcosms assessed through differential abundance analysis (DESeq2). 5-fold and higher differences are show and significant differences are denoted (*).

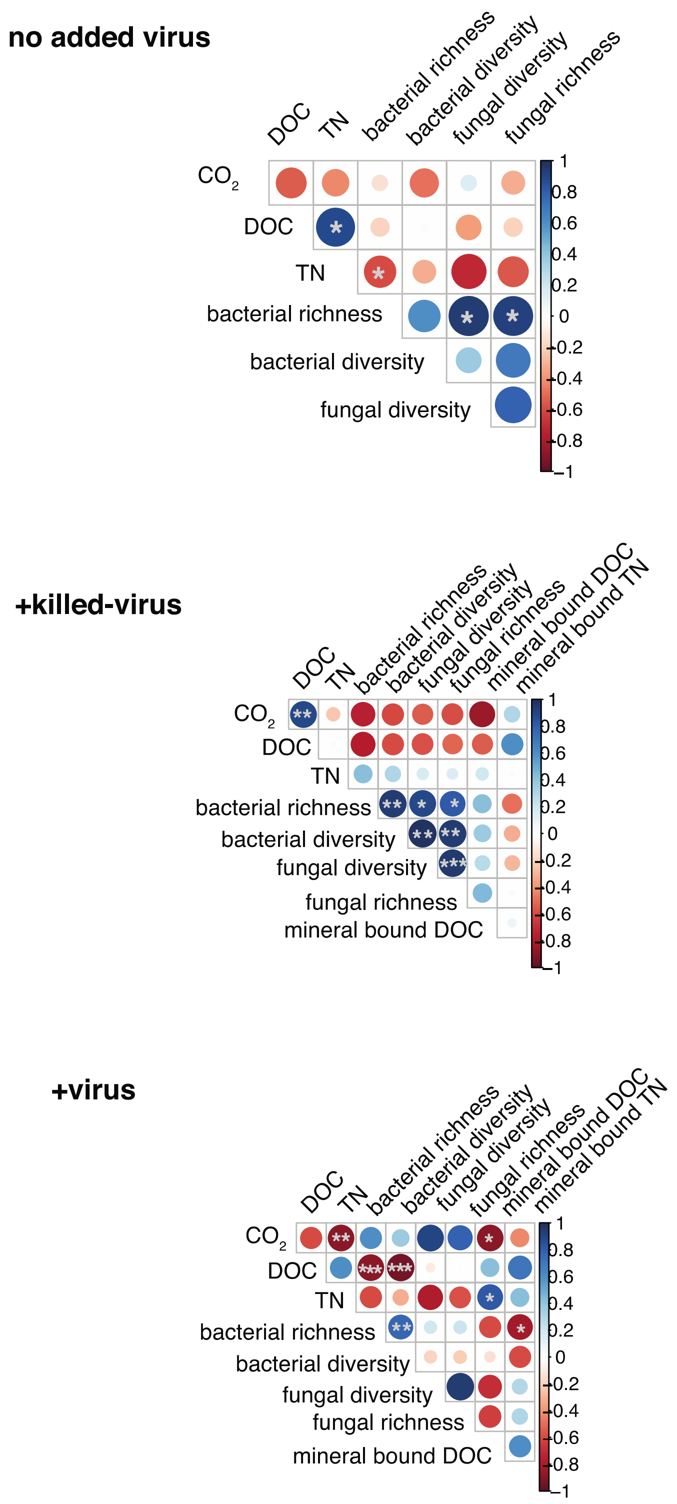


**Figure S7.** Correlations between carbon, nutrient, and microbial community traits across the three treatments for both soils combined. All spearman correlations are shown; correlations with p-values <0.05, 0.01, and 0.001 are denoted with a *,**, and *** respectively.
