## Supplemental Methods for "Experimental evidence for the impact of soil viruses on carbon cycling during surface plant litter decomposition"

*Soil collection, whole microbial community inoculum, and virus extraction*

Soils were collected from two locations, Pajarito Mountain Los Alamos (PJ) (35.894208° N, 106.391817° W) and Mount Baldy Santa Fe (SF) (35.793527° N, 105.800391° W) in northern New Mexico. These sites are at elevations are 2853m and 2952m respectively and dominated by pondersa pine and grasses, however, soil samples were collected from areas or bare soil away from immediate vegetation. Soils were stored for up to one month at 4°C. Before use soils were sieved with a 2mm sieve.

For each soil type we used 1 gram of the previously sieved and homogenized soil to create a whole community microbial inoculum. To make the microbial inoculum (50x soil dilution) we added 1 gram of soil to 9 mL of M9 media, shook the soil suspension and then let it sit for 2 minutes to allow particles to settle particles. We then added 1 mL of the suspension to 4 mL of M9 Media.

To concentrate and extract a viral fraction from each soil we used a combination of centrifugation and size filtering to exclude larger microbes. We extracted viruses from 300 grams of the two soil types. 50 grams of soil was added to 10 mL of M9 Medium and incubated overnight at 37°C. This was replicated 6 times for each soil. After the overnight incubation, we further split each sample to allow for centrifugation in following steps. 22 grams of soil slurry (soil and M9 Medium) was placed into twelve 50 mL falcon tubes. We added 26 mL of Glycine buffer (0.25N pH 9.5 – Tween 80 0.02% v/v) to each tube and incubated them at room temperature (25°C) for one hour while rotating. Samples were centrifuged at 4500 g for 15 minutes at 4°C and the supernatant was collected. The supernatant was then filtered using a 0.22 uM PVDF syringe Millipore filter. After filtering, we used Amicon Ultra filters to concentrate the sample. First, 50ml Amicon Ultra filters were coated with M9 buffer to reduce virus adherence to the membrane. This was achieved by adding 1 mL of M9 buffer to basket, pipetting up and down and then removing the M9 liquid. Next, we added 15 mL of our filtered samples and centrifuged at 3000 rpm for 20 minutes (25 C). Liquid flow through was discarded and we repeated this step until all of the sample was added to the Amicon filter and approximately 200-400 uL of liquid (concentrated virus) remained in the upper part of the tube. We add 100 uL of M9 media and pipette up and down to mix and resuspend the virus from the Amicon filter. Overall, after pooling samples we obtained 8 mL of viral concentrate per initial 300 grams of soil. An aliquot of virus concentrate from each soil was autoclaved to create a “killed-virus” sample.

In order to test the efficiency of our virus extraction technique, we created control samples for each soil where lambda phage (ATCC 97537) was spiked into a soil slurry (soil and M9 Medium) and extracted using the method outlined above. For these controls soil slurries were not incubated overnight. Following the phage extraction, we performed plaque assays to assess extraction efficiency. Lambda phage susceptible *E.coli* (ATCC 47076) were grown overnight in lambda broth mixing on a rotator at 37°C. We added 0.3 ml of *E.coli* and 0.1 mL of phage sample and mixed in a 2 mL tube. Phage samples included extractions from the two lambda spiked soils, a control of the initial lambda phage that was spiked into soils, and lambda phage that had been autoclaved to kill the phage. Serial dilutions of each sample type was tested and an *E.coli* control with no phage addition was included. After mixing the *E.coli* and phage, we incubated tubes at room temperature (25°C) for 20 minutes and then moved tubes to a heat-block where they were incubated at 37°C for 10 minutes. 0.4 mL of *E.coli* and phage mix was then added to 2.5 mL of heated lambda top agar and poured on pre-prepared lambda plates.

Plates were placed in the incubator at 37°C overnight and plaques were counted on each plate and used to compute the concentration of viable phage in each sample.

*Microcosm experiment*

Microcosms were constructed in 125 mL serum bottles with approximately 7 g of sand and 0.1 g (dry weight) of *Blue grama* grass litter cut into 1 cm pieces. The microcosms were sterilized by autoclaving (at 121°C and 15 psi) three times for 1 h each, with at least a 12-h resting interval between each autoclave cycle.

For each soil type (n=2) we had three different treatments, these included 1) whole microbial community (no added virus), 2) whole microbial community and killed virus extraction (+ killed-virus), and 3) whole microbial community and virus extraction (+virus) (Figure 1) with three replicates per treatment type for a total of 18 microcosms. The whole microbial community and virus extractions treatments were always from the same soil. We first created a homogenized inoculum for each treatment by adding viral extract to whole microbial inoculum at a ratio of 2:1. For the no added virus treatment we added M9 Media. We then add 1.5 mL of homogenized inoculum to each sterilized grass litter and sand microcosm. Microcosms were sealed with Teflon-lined crimp caps (preventing desiccation) and incubated at 25°C with a 12-hour light-dark cycle for 41 days. On days 2, 5, 8, 14, 21, 32, and 41, $CO_2$ was measured by gas chromatography using an Agilent Technologies 490 Micro GC (Santa Clara, CA, United States). After each measurement, the headspace air was evacuated with a vacuum pump and replaced with sterile-filtered air.

After the 41-day incubation, microcosms were destructively sampled to measure DOC, TN, and community composition. For each microcosm, 7 ml of sterile deionized water was added, swirled gently by hand for 30 s and then filtered through a 0.2 µm filter. To assess variation in DOC/TN composition or quality, the fraction of organic matter able to bind to mineral surfaces was measured on an aliquot of all samples following the protocol outlined in (1). DOC and TN concentration of each sample was measured on an OI Analytical model 1010 wet oxidation TOC analyzer (Xylem Inc., Rye Brook, NJ, United States). Following DOC/TN sampling, sand and litter material from each microcosm was frozen at −80°C for DNA extraction.

*DOC binding assay*

Aluminum oxide was used as a representative mineral (2) to assess DOC binding potential for the +killed-virus and +virus treatments as well as the initial viral concentrate used to inoculate these treatments. For each microcosm sample, 0.5 ml of DOM was added to 1.00 ml of sterile water (3X dilution factor) and 0.3 grams of aluminum oxide ($Al_2O_3$). A 6X dilution was used for inoculum samples. Samples were mixed by inversion with a Multi-Purpose Rotator Model 151, speed 4 (Scientific Industries Inc, NY) for 30 minutes and then centrifuged at 16,100 x *g* for 5 minutes. Supernatant was transferred to a new tube and stored at -20°C until DOC quantification on a TOC analyzer. The percentage of bound DOC was calculated as 100% x (1-($DOC_{post\text{-}binding}$ x dilutionFactor)/$DOC_{pre\text{-}binding}$).

*Microbial community taxonomic profiling*

For the final microcosm samples, we extracted and sequenced DNA to obtain bacterial (16S rRNA) and fungal (ITS) community profiles. DNA extractions were performed with a DNeasy PowerSoil Kit (Qiagen, Hilden, Germany) following the manufactures protocol with the following exceptions, 0.3g of material was used per sample extract and all samples were eluted to a final volume of 30ul. DNA samples were quantified with the Invitrogen Qubit dsDNA HS Assay Kit (Invitrogen,Waltham, MA), on an Invitrogen Qubit 2.0 following the manufactures protocols.

The bacterial 16S rRNA genes were amplified with methods previously described in (1) with the following modifications. Following step one of PCR the PCR products were cleaned with a 0.9 Ratio of Beckman Coulter Agentcourt AMPure XP beads (Beckman Coulter, Brea, CA) followed by step two of PCR. Amplicons were cleaned using the same method as the PCR1 products and quantified using the same procedure as the extracted DNA, then pooled to 10ng each. The pool was then cleaned with beads in the same manner as above following manufactures protocol. The fungal ITS regions were amplified using an equimolar mixture of three ITS9 forward primers (ITS9f_FS1: TCGTCGGCAGCGTCAGATGTGTATAAGAGACAGNNNNNNGAACGCAGCRAAIIGYGA, ITS9f_FS2: TCGTCGGCAGCGTCAGATGTGTATAAGAGACAGNNNNNGAACGCAGCRAAIIGYGA, and ITS9F_FS3: TCGTCGGCAGCGTCAGATGTGTATAAGAGACAGNNNNGAACGCAGCRAAIIGYGA) and the ITS4r_FS reverse primer (GTCTCGTGGGCTCGGAGATGTGTATAAGAGACAGNNNNNNTCCTCCGCTTATTGATATGC) (3). Amplification procedure was used based on Gloor et al. 2010, with Phusion Hot Start II High Fidelity DNA polymerase (Thermo Fisher Scientific, Vilnius, Lithuania). In the first PCR, barcoded amplicons were produced over 25 cycles using gene primers flanked by 6 nt barcodes that jointly provided a unique 12-mer barcode for each sample (4). Cycling conditions were 5 min at 95°C, 25 cycles of (95°C for 30 s, 50°C for 60 s, 68°C for 60 s), and a final extension step of 68°C for 10min. The second PCR extended Illumina adapter sequences on the amplicons over 12 cycles. Cycling conditions were 5 min at 95°C, 12 cycles of (95°C for 30 s, 50°C for 60 s, 68°C for 60 s), and a final extension step of 68°C for 10min. Amplicons were cleaned and concentrated using the same procedure as with the bacterial 16S rRNA genes described above. DNA quality of the bacterial and fungal amplicon pools were assessed with a bioanalyzer, concentration was verified by qPCR, and sequencing was performed on an Illumina MiSeq with paired-end 250 bp chemistry at Los Alamos National Laboratory. Unprocessed sequence data are available through NCBI’s Sequence Read Archive (PRJNA763874)

Sequence data were processed using UPARSE (5), where the same methods as previously described in (6) where used to obtain OTU tables. OTU tables were rarefied to the lowest number of common sequences for bacterial and fungal profiles (n=1560 and n=2043, respectively). OTU tables were used to calculate Bray-Curtis distance matrices and diversity metrics (richness and Shannon diversity).

*Statistical analysis*

Many of the statistical tests were performed separately for each microbial inoculum (PJ and SF) as we expected substantial differences in the microbial composition and nutrient inputs across the different inocula due to the geographic distance of their initial collection points. All statistical analyses and graphing were performed in the R software environment. To test for differences in function among treatments we performed ANOVA analyses (aov function, R). More specifically, we tested the impacts of treatment on $CO_2$ production over time and DOC and TN at 41 days in SF and PJ microcosms. In addition, we tested the impacts of treatment on mineral binding of DOC and TN. To test for differences in bacterial and fungal composition among treatments we performed PERMANOVA analyses (vegan package R). To identify taxa contributing to significant compositional differences, we used indicator species analysis (7). We also performed an alternative where we identified differentially abundant taxa between the +killed-virus and +virus treatment using a DESeq2 analysis (8). Lastly, we combined data across both soils and for each treatment we tested for correlations between bacterial and fungal richness/diversity and carbon and nutrient measurements using the spearman method.
